# ClearGel: a thermoresponsive hydrogel for chronic, near-millimeter two-photon imaging through the intact skull

**DOI:** 10.64898/2026.11.04.764111

**Authors:** Lina Liu, Yufeng Gao, Yuezhi He, Na Xiao, Jia Yu, Ting Wu, Shiwei Ye, Jiuling Liao, Liang Wang, Jun Chu, Liangjian Liu, Chengbo Liu, Xuemin Du, Hui Li, Wei Zheng

## Abstract

The intact skull remains a fundamental barrier to high-resolution two-photon neuroimaging in live mice. Existing skull-clearing methods, however, cannot simultaneously provide deep cortical access, spatial selectivity, reversibility, and biological safety. Here we introduce ClearGel, a thermoresponsive hydrogel that undergoes rapid *in situ* gelation upon contact with the skull, confining the clearing agents to the targeted region while sustaining their release over time. ClearGel routinely enables two-photon imaging down to approximately 800 μm through the intact skull—a depth statistically equivalent to open-skull windows and substantially surpassing all prior approaches. It supports user-defined clearing patterns from focal sites to whole-skull coverage, allows repeated imaging over 14 days through a reversible optical interface, and, critically, enables awake functional imaging of both neuronal Ca^2+^ activity and the more elusive cAMP dynamics at single-cell resolution. ClearGel requires only simple topical application—no craniotomy or skull thinning—and provides a minimally invasive, chronic-compatible optical window for longitudinal studies of cortical function in behaving mice.

## Introduction

Two-photon microscopy (TPM), in conjunction with genetically encoded indicators, has become a cornerstone modality for *in vivo* mouse brain imaging, offering high-resolution structural and functional readouts at exceptional penetration depths^1-3^. However, the intrinsic optical scattering and aberration imposed by the skull severely compromise image quality during transcranial TPM^4-6^. To circumvent this barrier, open-skull cranial windows and thinned-skull preparations have been widely adopted^5,7,8^. Open-skull windows afford superior optical access and support chronic imaging, yet they are inherently invasive, disrupting the native physiological milieu through tissue injury, glial inflammatory responses, and cerebrospinal fluid loss^5,9^. Thinned-skull approaches are less invasive but remain technically demanding, particularly in achieving and maintaining a uniformly thin bone layer for longitudinal experiments^7,8^. Hence, a minimally invasive alternative that preserves skull integrity while enabling high-quality cortical imaging is highly desirable.

Skull optical clearing, which preserves brain physiological integrity and enables non-invasive transcranial access, has emerged as a compelling solution^6,10-14^. Over the past decade, liquid-agent-based methods have made notable strides, but their clearing efficacy is fundamentally constrained by short incubation times (typically 5–35 min *in vivo*), limiting imaging depths to approximately 150–400 μm (e.g., SOCW^12^, USOCA^10^, SeeThrough^6^). UV-curable technologies (LCCW^11^, TIS^14^) were introduced to address reagent loss by forming a solid film upon UV irradiation, extending depth to 400–500 μm, yet they cannot replenish agents post-curing, causing gradual quality decline. Our previously developed HOTS strategy employed a head-mounted device that enabled clearing in awake, freely moving mice, extending incubation time and achieving TPM visualization down to ∼700 μm^13^. However, this depth capability came at the cost of a bulky, surgically affixed apparatus that limited experimental throughput, and the prolonged direct contact of high-concentration liquid reagents raised concerns regarding tissue irritation and potential inflammation. Despite these advances, no existing method simultaneously offers (i) deep cortical imaging comparable to open-skull windows, (ii) spatial selectivity without physical barriers, (iii) a reversible interface for chronic re-imaging, and (iv) minimal biological perturbation—a combination of features essential for routine, high-throughput longitudinal neuroscience studies.

Here we introduce ClearGel, a thermoresponsive hydrogel-based skull optical clearing agent for intact skull brain imaging. At room temperature, ClearGel exists as a low-viscosity sol for straightforward application; upon contact with the warm skull (∼37 °C), it rapidly undergoes *in situ* gelation, forming a stable, localized, and conformal transparent layer. This temperature-triggered phase transition transforms the clearing process from a freely diffusive liquid treatment into a gel-confined surface reaction. Notably, the hydrogel matrix itself exhibits an intrinsic optical clearing effect, while its sustained-release property maintains a prolonged concentration gradient across the full skull thickness. These two effects, together with the optimized clearing agent formulation, achieve more thorough skull clearing, enabling routine TPM down to ∼800 μm through the intact skull with image quality comparable to that of open-skull windows. Crucially, unlike HOTS, ClearGel accomplishes this without any head-mounted hardware, and its gel-based sustained release inherently avoids the osmotic and chemical shock associated with direct liquid flooding, thereby minimizing tissue irritation and glial activation. In addition, because the treated region is precisely delineated by the gel-covered area, ClearGel permits spatially separated multi-site clearing, offering flexible selection from local spots to whole-skull coverage without physical barriers—a feature particularly valuable for intra-animal contralateral comparisons and multi-region longitudinal tracking. Unlike UV-cured windows that permanently seal the skull and preclude reagent replenishment, ClearGel’s reversible format allows re-establishment of the optical interface before each imaging session. ClearGel is the first skull-clearing strategy that simultaneously achieves open-skull-comparable imaging depth, reversible chronic access, and spatially selective clearing without physical barriers. We anticipate that this hydrogel-based strategy will serve as a practical, high-performance alternative to invasive craniotomy, facilitating routine high-throughput longitudinal studies of cortical plasticity, vascular dynamics, and neuroimmune interactions in awake, behaving mice.

## Results

### Development and properties of ClearGel

To address the challenges of uncontrolled spreading and poor spatial confinement of freely diffusible liquid optical clearing agents (OCAs) on the curved skull surface, we sought to identify a hydrogel system capable of storing and sustained release of the clearing reagents. Because conventional hydrogels may leave interfacial gaps upon contact with the skull, compromising clearing efficacy, we focused on thermoresponsive hydrogels that undergo sol–gel transition under physiological conditions (Fig. 1a, b)^15-17^. We first screened natural and synthetic hydrogel-forming polymers based on three criteria: biocompatibility, simple preparation, and ease of handling. Ten candidates were selected for initial evaluation: agar, gelatin, sodium alginate/calcium chloride, carrageenan, P188, carbopol 940, HPMC, chitosan/β-glycerophosphate (chitosan/β-GP), P407, and PLGA–PEG–PLGA. Among these, only chitosan/β-GP, P407, and PLGA–PEG–PLGA exhibited thermoresponsive sol–gel transition properties at physiological temperature (∼37 °C) and were therefore selected for further evaluation (Fig. 1e).

**Fig 1.**
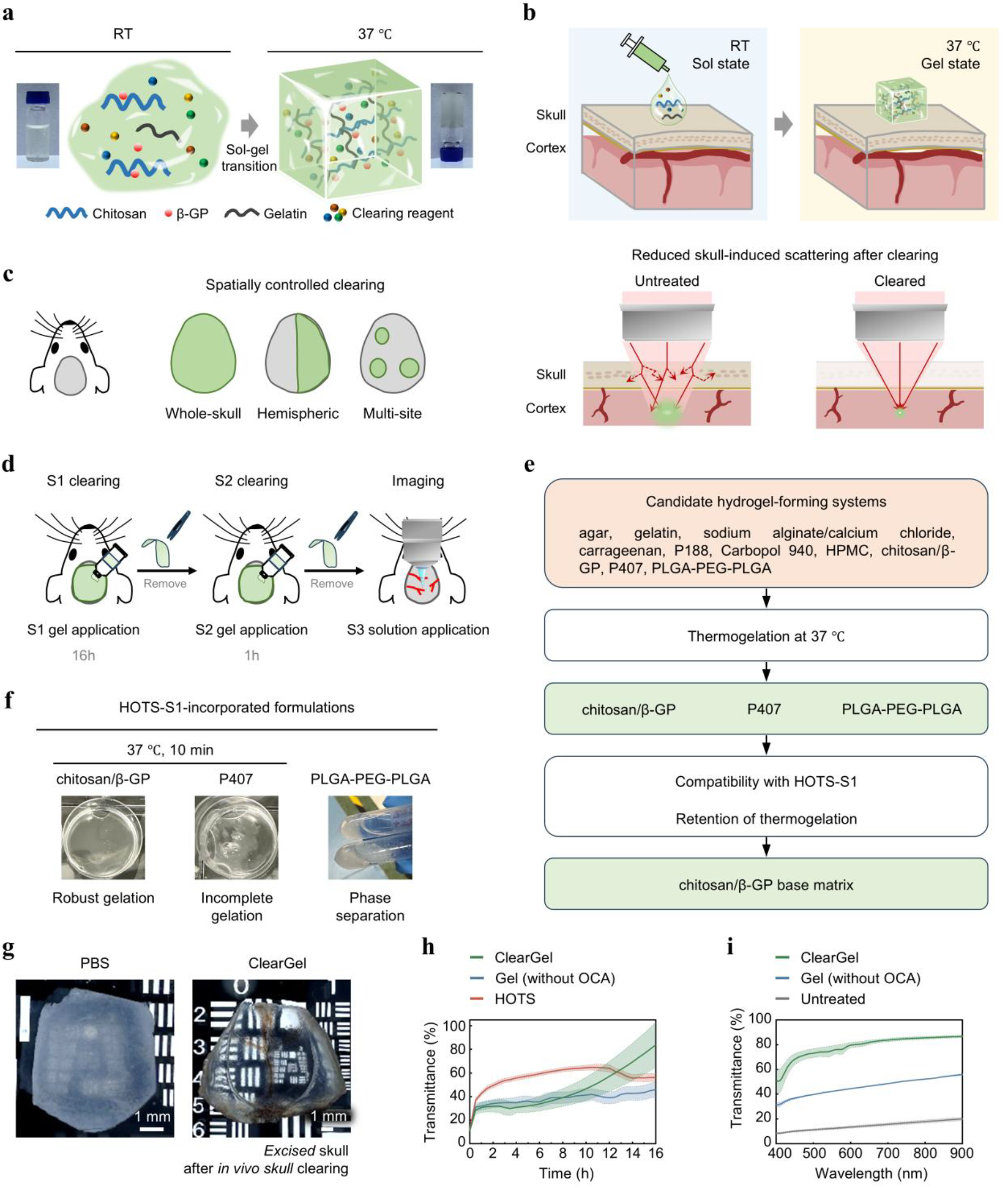
ClearGel enables thermoresponsive and spatially confined skull optical clearing. **a**, Schematic and representative photographs showing the temperature-triggered sol–gel transition of ClearGel. ClearGel, composed of chitosan, β-glycerophosphate (β-GP), gelatin and optical clearing agents (OCAs), remains flowable at room temperature (RT) and forms a stable hydrogel at 37 °C. **b**, Schematic of ClearGel application and *in situ* gelation on the skull surface. Sol-state ClearGel is applied at RT and gels upon contact with the warm skull, retaining the clearing formulation over the treated region. Bottom, schematic illustrating the reduction in skull-induced light scattering after clearing. **c**, Schematic of whole-skull, hemispheric, and spatially separated multi-site clearing patterns achieved by varying the ClearGel-covered area. **d**, Sequential *in vivo* ClearGel treatment and imaging workflow. ClearGel-S1 was applied for 16 h and removed, followed by ClearGel-S2 for 1 h. After removal of ClearGel-S2, a hydrogel-free S3 refractive-index matching solution (RI = 1.47) was applied as the imaging medium for two-photon microscopy. **e**, Screening workflow used to identify the thermoresponsive hydrogel matrix. Ten candidate hydrogel-forming systems were screened for thermogelation at 37 °C. Chitosan/β-GP, P407 and PLGA-PEG-PLGA exhibited thermoresponsive gelation and were further evaluated for compatibility with HOTS-S1 and for retention of thermogelation after HOTS-S1 incorporation. Chitosan/β-GP was selected as the base matrix for ClearGel. **f**, Representative photographs of chitosan/β-GP, P407 and PLGA-PEG-PLGA formulations after HOTS-S1 incorporation and incubation at 37 °C for 10 min. Chitosan/β-GP retained robust thermogelation, whereas P407 formed an incomplete gel and PLGA–PEG– PLGA underwent phase separation. **g**, Representative transmitted-light images of excised skulls from 6-week-old C57BL/6J mice following the *in vivo* clearing procedure. **h**, Time course of skull transmittance at 600 nm during *ex vivo* treatment with ClearGel, Gel, and HOTS over 16 h. ClearGel was allowed to gel at 37 °C for 3 min before room temperature clearing. Gel is the chitosan/β-GP/gelatin hydrogel matrix alone without optical clearing agents. Data are mean ± s.e.m.; n ≥ 2 mice. **i**, Spectral transmittance of excised skulls from 6-week-old C57BL/6J mice following *ex vivo* treatment with ClearGel, PBS or Gel (chitosan/β-GP/gelatin). Data are mean ± s.e.m.; n ≥ 2 mice.

We next assessed the drug-loading capacity of these candidates. The optical clearing formulation of ClearGel is derived from our previously developed HOTS strategy^13^; we therefore examined whether the three thermoresponsive hydrogels were compatible with the HOTS OCAs. In the HOTS protocol, the clearing agents are divided into HOTS-S1 and HOTS-S2. Because EDTA-mediated decalcification is a critical step in skull clearing, and HOTS-S1 contains the complete set of chemical species used in the HOTS clearing protocol—including EDTA—we selected EDTA alone and complete HOTS-S1 for compatibility testing. Specifically, we evaluated whether each hydrogel system remained homogeneous and retained gelation at physiological temperature after loading with EDTA or HOTS-S1. PLGA–PEG–PLGA underwent phase separation after mixing with either EDTA or HOTS-S1. P407 remained homogeneous after mixing with EDTA or HOTS-S1 but formed only a weak and partial gel at 37 °C after loading with HOTS-S1. In contrast, chitosan/β-GP remained homogeneous after mixing with either EDTA or HOTS-S1 and retained stable thermoresponsive gelation at 37 °C after HOTS-S1 loading (Fig. 1f). We therefore selected chitosan/β-GP as the base hydrogel system.

Having identified a suitable thermoresponsive hydrogel, we then optimized the clearing reagent composition. The core principle was to maximize clearing efficacy while preserving the thermoresponsive gelation properties. We also balanced gel stability, handling properties, and the kinetics of the sol–gel transition^17-20^. This optimization yielded two thermoresponsive hydrogel-based optical clearing reagents, termed ClearGel-S1 and ClearGel-S2, formulated to retain the sequential clearing functions of HOTS-S1 and HOTS-S2 while converting the liquid skull clearing process into a gel-confined procedure (detailed formulations are provided in Methods). ClearGel-S1 was designed primarily for decalcification, collagen dissociation, and delipidation, whereas ClearGel-S2 was designed for refractive-index (RI) matching, further collagen dissociation, and skull homogenization. The two reagents were sequentially applied to the skull surface and removed before imaging. After ClearGel-S2 removal, a hydrogel-free RI-matching S3 solution (RI: 1.47), based on the ClearGel-S2 recipe, was applied as the imaging medium for TPM (Fig. 1d). Because ClearGel-S1 and ClearGel-S2 undergo *in situ* gelation directly on the warm skull surface, this workflow enables gel-confined skull optical clearing without requiring a chamber or cap. Consequently, the cleared region is defined by the gel-covered area, simplifying the clearing procedure and enabling spatially controllable skull clearing (Fig. 1c; Extended Data Fig. 1; Supplementary Video 1 and Video 2).

We then evaluated the optical clearing performance of ClearGel using *ex vivo* skulls harvested after the *in vivo* clearing workflow. Bright-field imaging showed that ClearGel markedly increased skull transparency compared with PBS-treated samples (Fig. 1g). Quantitative measurements further revealed that ClearGel increased skull transmittance from 13% to 80% (Fig. 1i). To probe the underlying mechanism, we next compared the clearing kinetics of ClearGel and HOTS using paired *ex vivo* measurements of contralateral hemiskulls. As a control, we also monitored the dynamic effect of the hydrogel matrix alone (without OCA) on isolated skulls. As shown in Fig. 1h, HOTS induced a rapid initial increase in transmittance, reached a plateau at approximately 3 h, increased only modestly thereafter, and declined at approximately 12 h. In contrast, ClearGel exhibited a slower onset of clearing, with transparency gradually accelerating after approximately 10 h, reaching nearly complete transparency at around 16 h, and ultimately surpassing the final clearing effect of HOTS (Fig. 1h). Furthermore, we found that the hydrogel matrix alone also exhibited a certain degree of clearing effect, which increased over time; after 16 h of treatment with the hydrogel matrix alone, skull transmittance increased from 13% to 45%. These results suggest that the superior clearing performance of ClearGel arises from the intrinsic clearing effect of the hydrogel matrix combined with its sustained-release property. The combination of these two effects, together with the optimized clearing agent formulation, enables ClearGel to outperform nearly all previously reported *in vivo* skull clearing strategies.

### ClearGel enables deep transcranial two-photon imaging through the intact skull

To quantify the imaging performance enabled by this clearing strategy, we first compared ClearGel with representative skull-retaining preparations, including HOTS^13^, SeeThrough^6^, and the widely used thinned-skull preparation^7,8^. Under identical imaging conditions, ClearGel consistently enabled superior penetration depth and neuronal structural contrast compared with all three reference preparations (Fig. 2a–b). In 6-week-old Thy1-GFP mice, the maximum imaging depth through the untreated skull was less than 200 μm below the pia. After ClearGel clearing, the average imaging depth increased to approximately 800 μm, outperforming SeeThrough (∼400 μm), thinned-skull (∼500 μm), and HOTS (∼700 μm) (Fig. 2b). Depth-resolved maximum intensity projection (MIP) images revealed that, despite expected degradation at deeper layers, ClearGel maintained discernible neuronal somata at 700–870 μm below the pia, spanning deep layer 5 and the putative upper layer 6 region—a depth at which SeeThrough failed to resolve individual cell bodies (Fig. 2c). Notably, although the clearing reagent formulation of ClearGel is optimized from that of HOTS, the intrinsic clearing effect of the hydrogel matrix combined with its sustained-release property enhances clearing homogeneity and depth beyond what the liquid-based HOTS can achieve. *In vivo* second harmonic generation (SHG) imaging further demonstrated that ClearGel dissociated skull collagen more completely than HOTS (Extended Data Fig. 2). Collectively, these results establish ClearGel as a superior skull-retaining clearing strategy, markedly outperforming existing liquid-based methods and the thinned-skull preparation in terms of penetration depth and image quality.

**Fig 2.**
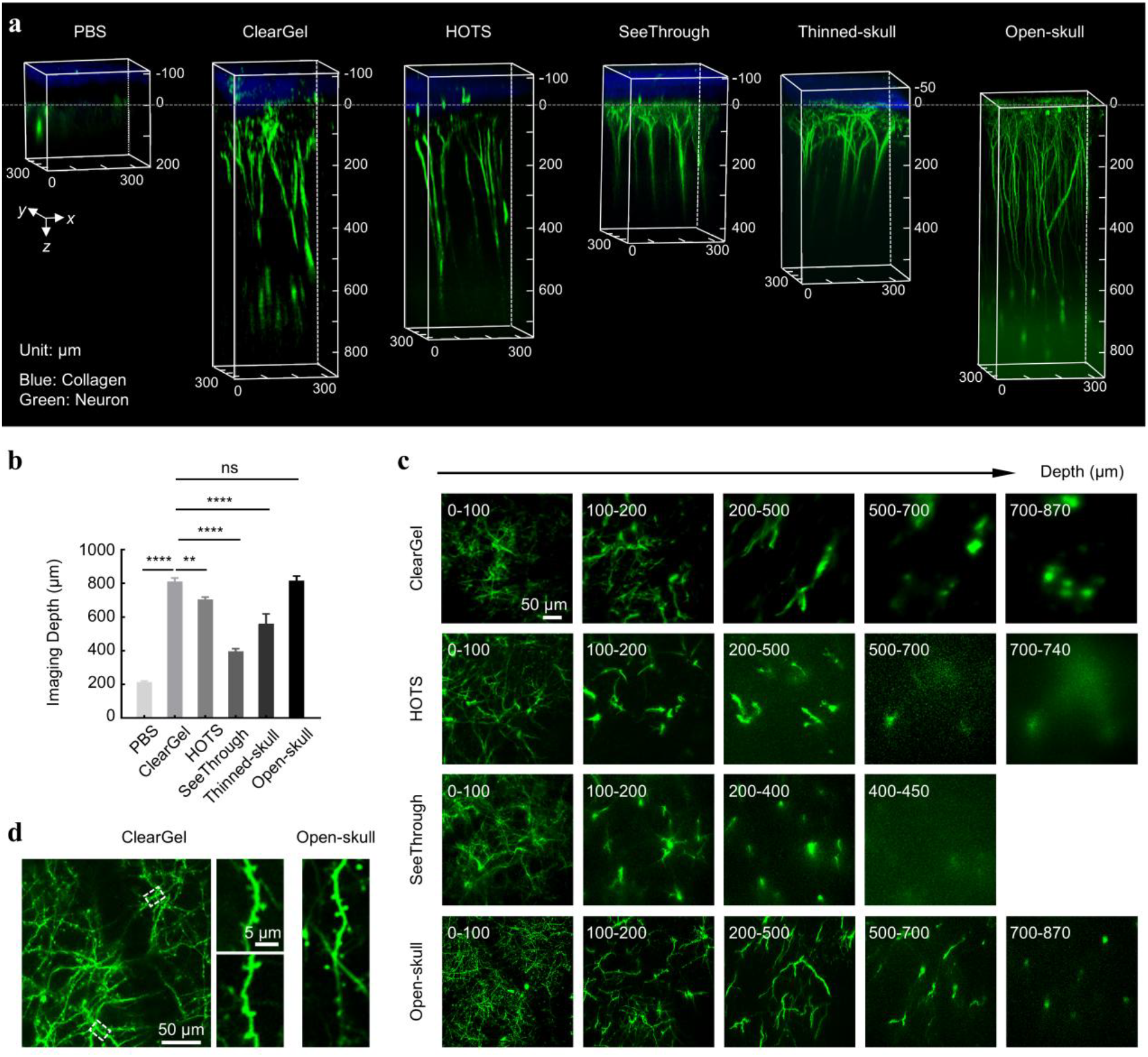
ClearGel supports deep neuronal imaging through the intact skull at depths approaching those achieved through open-skull windows. **a**, Representative three-dimensional *in vivo* two-photon reconstructions of EGFP-expressing cortical neurons acquired in 6-week-old Thy1-GFP-M mice under the indicated cranial conditions. Green, neuronal EGFP fluorescence; blue, second-harmonic generation (SHG) signal from skull collagen. Depth is referenced to the pial surface, defined as 0 µm; negative values indicate positions above the pia. **b**, Quantification of maximum imaging depth, defined as the distance from the pial surface to the deepest imaging plane at which neuronal structure could still be reliably identified. Data are mean ± s.e.m.; n ≥ 3 mice. Statistical significance was assessed using one-way ANOVA followed by Tukey’s multiple-comparisons test. Only comparisons with ClearGel are shown. NS, not significant (P > 0.05); **P < 0.01, ****P < 0.0001. **c**, Representative depth-resolved maximum-intensity projections of EGFP-expressing neuronal dendrites and somata acquired under the indicated cranial conditions. Numbers indicate depth ranges relative to the pial surface. **d**, Representative two-photon images of cortical dendrites and dendritic spines acquired through a ClearGel-treated intact skull and an open-skull window. For ClearGel, enlarged views of the regions indicated by the white dashed boxes are shown.

We next benchmarked ClearGel against open-skull cranial windows, which physically remove the skull and thus provide the gold standard for transcranial imaging quality^5^. Through ClearGel-cleared skulls, we readily resolved neuronal somata in layer 5 at depths comparable to those achieved through open-skull windows (Fig. 2a and Supplementary Video 3). While the open-skull preparation yielded slightly superior resolution at extreme depths—primarily attributable to spherical aberrations arising from refractive index mismatch between the clearing medium and brain tissue, which progressively degrade image quality with increasing depth—ClearGel performed remarkably well in superficial layers. At the level of dendritic spines, ClearGel achieved resolution and contrast that closely approached those of open-skull windows, with clearly resolved spine morphology (Fig. 2d). Thus, while deep-layer resolution remains modestly inferior to the open-skull gold standard, ClearGel delivers open-skull-comparable imaging performance in the superficial-to-intermediate cortical layers while fully preserving skull integrity and avoiding the physiological disruptions associated with craniotomy.

### ClearGel facilitates transcranial imaging of neuronal Ca^**2+**^ and cAMP dynamics

We next asked whether the optical access provided by ClearGel was sufficient for functional imaging through the intact skull. Neuronal Ca^2+^ transients are widely used to monitor cortical activity^21,22^, whereas intracellular second-messenger dynamics can further reveal regulatory signaling underlying neural circuit function^23,24^. We therefore first imaged stimulus-evoked neuronal Ca^2+^ activity through the ClearGel-treated skull. In Thy1-GCaMP6s mice, hindpaw electrical stimulation evoked robust Ca^2+^ transients in neurons of the somatosensory cortex (Fig. 3a–e and Supplementary Video 4). Images acquired before, during, and after stimulation revealed clear fluorescence increases in multiple neuronal somata (Fig. 3b). Single-cell traces showed rapid, stimulus-aligned Ca^2+^ responses (Fig. 3c). ΔF/F_0_ response maps delineated the spatial distribution of responsive cells across the field of view (Fig. 3d), and repeated stimulation produced reproducible response patterns across trials (Fig. 3e). These results demonstrate that ClearGel supports transcranial imaging of stimulus-evoked neuronal Ca^2+^ activity at single-cell resolution in awake mice.

**Fig 3.**
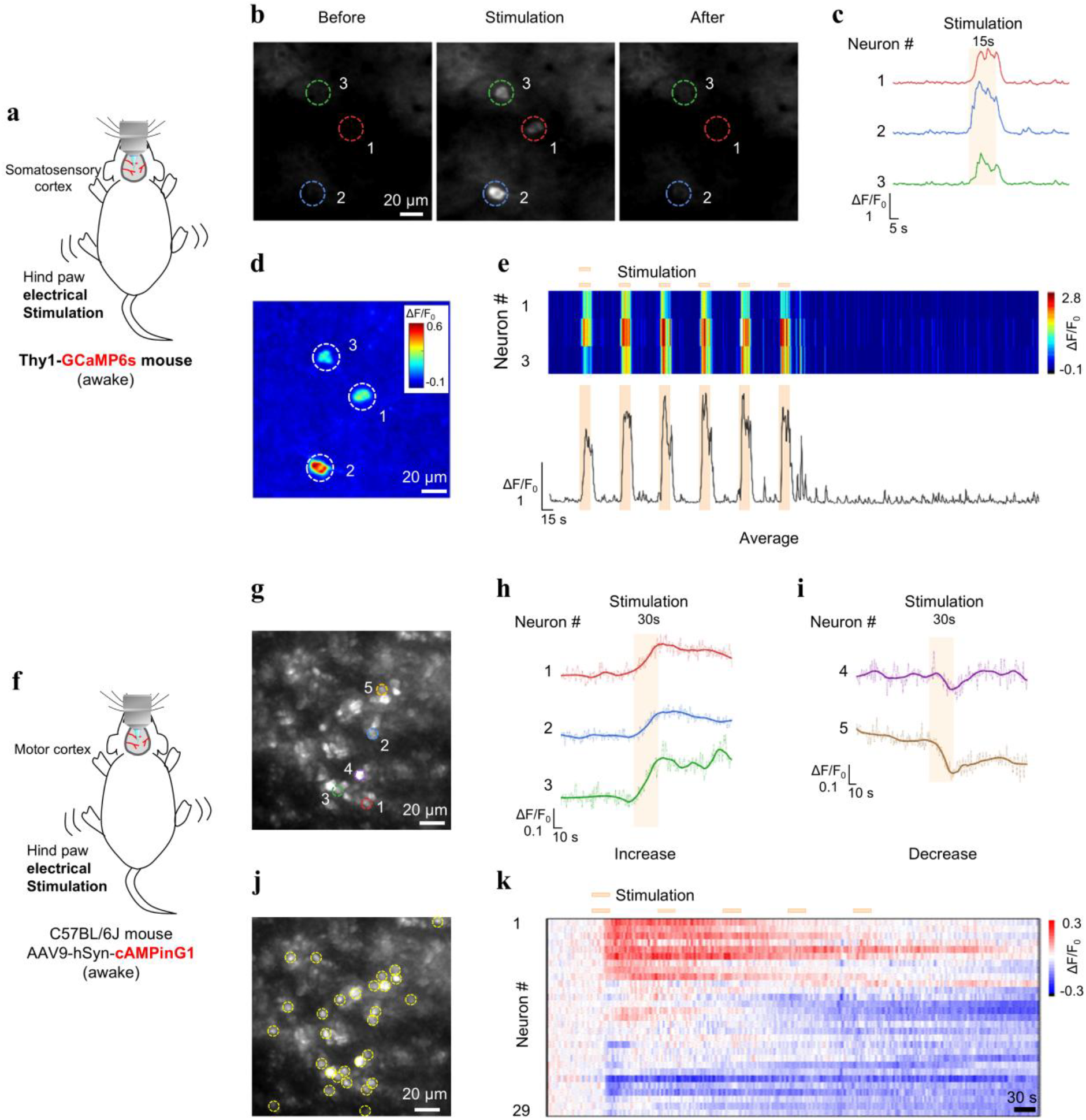
ClearGel enables transcranial imaging of neuronal Ca^**2+**^ and cAMP dynamics at cellular resolution in awake mice. **a**, Experimental schematic for two-photon Ca^2+^ imaging in the somatosensory cortex of awake 6-week-old Thy1-GCaMP6s mice through the ClearGel-treated intact skull. Imaging was performed at 200 µm below the pia during hind paw electrical stimulation (0.2 mA, 5 Hz, biphasic pulses; 15 s per trial). **b**, Representative images of the same field of view acquired before, during, and after stimulation. Dashed circles indicate the neurons analyzed in **c. c**, ΔF/F_0_ traces from the three neurons indicated in **b** during a representative stimulation trial. The shaded region indicates the stimulation period. **d**, ΔF/F_0_ response map for the field of view shown in **b**. Dashed circles indicate the same neurons. **e**, Heat map of ΔF/F_0_ responses from the three neurons during six repeated stimulation trials. The trace below shows the mean ΔF/F_0_ across the three neurons. Orange bars indicate stimulation periods. **f**, Experimental schematic for two-photon cAMP imaging in the primary motor cortex of awake 8–9-week-old C57BL/6J mice expressing AAV9-hSyn-cAMPinG1 through the ClearGel-treated intact skull. Imaging was performed at 180 µm below the pia during hind paw electrical stimulation (0.2 mA, 5 Hz, biphasic pulses; 30 s per trial). **g**, Representative image of cAMPinG1-expressing cortical neurons. Colored circles indicate the neurons analyzed in **h** and **i. h, i**, ΔF/F_0_ traces from representative neurons exhibiting stimulus-associated increases (**h**) or decreases (**i**) in cAMPinG1 fluorescence. Light-colored traces show the raw ΔF/F_0_ signals, and dark-colored lines show the corresponding wavelet-denoised traces. Shaded regions indicate stimulation periods. **j**, Field of view shown in **g**, with 29 neuronal regions of interest used to generate the heat map in **k. k**, Heat map of raw ΔF/F_0_ traces from the 29 neurons during five repeated stimulation trials. Orange bars indicate stimulation periods. Panels **a**-**e** show one representative Thy1-GCaMP6s mouse; similar responses were observed in n ≥ 3 mice. Panels **f**-**k** show one representative cAMPinG1-expressing mouse; similar experiments were performed in n ≥ 3 mice.

We next asked whether ClearGel could resolve a more challenging functional signal—intracellular cAMP dynamics. cAMP is a key second messenger downstream of neuromodulatory GPCR pathways^23,24^, but its signals are generally weaker and more heterogeneous than Ca^2+^ transients^25^, rendering transcranial detection substantially more challenging. To test this, we imaged cAMP dynamics in awake C57BL/6J mice expressing the cAMP sensor cAMPinG1 in the motor cortex^24^. Hindpaw electrical stimulation induced clear stimulus-associated changes in cAMP fluorescence through the cleared skull (Fig. 3f–k and Supplementary Video 5). Notably, unlike the predominantly stimulus-aligned Ca^2+^ transients, individual neurons displayed heterogeneous cAMP response profiles, including both signal increases and decreases (Fig. 3h, i)—a pattern consistent with cell-specific regulation of cAMP signaling. Analysis of 29 neurons further revealed diverse, recurrent stimulus-modulated cAMP dynamics across repeated trials (Fig. 3j, k). Together, these results establish that ClearGel enables transcranial functional imaging of both neuronal Ca^2+^ activity and cAMP dynamics at single-cell resolution in awake mice, including detection of relatively weak and heterogeneous second-messenger signals.

### Spatially controllable optical access from whole-skull to selective regions

High-resolution optical imaging across the whole dorsal cortex remains a major challenge^2,26^. Craniotomy over large areas risks infection and tissue damage, while large-scale thinned-skull preparation is technically demanding and difficult to achieve uniformly^5,7-9^. Skull optical clearing offers a potential solution for whole-skull transparency; we therefore first examined whether ClearGel could clear the entire dorsal skull. Laser speckle contrast imaging revealed that following whole-skull ClearGel treatment, cortical vascular patterns became clearly visible across the dorsal surface, with substantially improved vessel contrast relative to untreated controls (Fig. 4a, b). Line profiles from representative regions showed marked increases in vascular signal over background after clearing, and quantification confirmed an increase in vessel signal-to-background ratio (Fig. 4d, e). Bright-field imaging further confirmed that the skull surface remained morphologically intact after treatment, with no visible bleeding at the cranial sutures (Fig. 4c). These results indicate that ClearGel provides broad optical access across the dorsal skull while preserving skull surface integrity.

**Fig 4.**
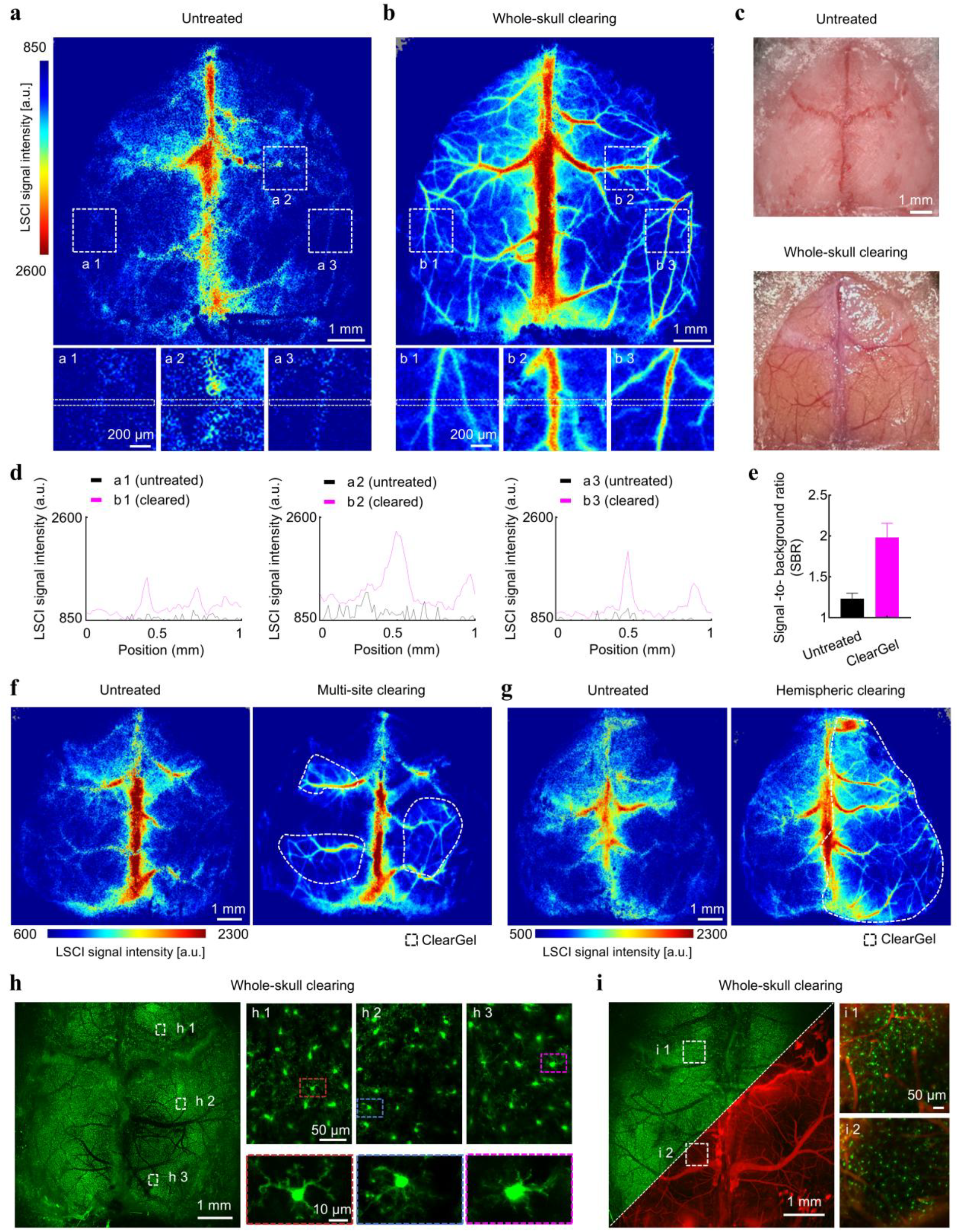
ClearGel enables spatially controlled optical access from focal cortical regions to the whole skull. **a, b**, Laser speckle contrast imaging (LSCI) maps of the dorsal skull from the same 6-week-old C57BL/6J mouse before (**a**) and after (**b**) whole-skull ClearGel treatment. Enlarged views of the boxed regions are shown below (**a1**–**a3** and **b1**–**b3**). **c**, Corresponding bright-field images of the dorsal skull from the same mouse acquired before and after whole-skull ClearGel treatment. **d**, LSCI signal-intensity profiles measured along the horizontal dashed line scans in the matched regions shown in **a1**-**a3** and **b1**-**b3**. Black, untreated; magenta, ClearGel-treated. **e**, Quantification of vascular signal-to-background ratio (SBR) calculated from four matched regions before and after ClearGel treatment; three representative regions are shown in **a** and **b**. Data are mean ± s.e.m.; *n* = 3 matched regions from the mouse shown in **a** and **b. f**, Representative LSCI maps of the dorsal skull from the same mouse before and after multi-site ClearGel treatment. Dashed outlines indicate the predefined target regions. **g**, Representative LSCI maps of the dorsal skull from the same mouse before and after hemispheric ClearGel treatment. The dashed outline indicates the treated hemisphere. **h**, Large-field-of-view two-photon maximum-intensity projection of EGFP-expressing cortical microglia acquired over a 6.45 × 6.45 mm^2^ field of view after whole-skull ClearGel treatment in a 6-week-old *Cx*_*3*_*cr1*^*GFP/+*^ mouse. Higher-magnification maximum-intensity projections of boxed regions **h1–h3**, acquired approximately 150-200 µm below the pia, are shown on the right. Bottom, enlarged views of the color matched boxed microglial cells. Example from n ≥ 3 mice. **i**, Large-field-of-view two-photon maximum-intensity projection of EGFP-expressing microglia and Texas Red dextran-labelled cortical vasculature after whole-skull ClearGel treatment in a 6-week-old *Cx*_*3*_*cr1*^*GFP/+*^ mouse. The overview is shown as a split-channel composite, with microglia in green and vasculature in red. Boxed regions **i1** and **i2** are shown as merged images at higher magnification on the right. Example from n ≥ 3 mice.

To verify that the cleared regions provided sufficient optical quality for cellular-resolution imaging across multiple areas of the dorsal cortex, we performed large-field-of-view transcranial two-photon microscopy after whole-skull clearing in *Cx*_*3*_*cr1*^*GFP/+*^ mice, in which microglia are natively fluorescent. ClearGel enabled wide-field mapping of cortical microglia across the dorsal cortex; higher-magnification views from selected regions resolved fine ramified microglial processes at cellular resolution (Fig. 4h; Extended Data Fig. 3; Supplementary Video 6). Imaging was performed at a depth of approximately 150– 200 μm below the pia, corresponding to cortical layer 2/3, where microglial somata and processes were clearly resolved. In mice additionally labeled with Texas Red dextran for vascular contrast, the same preparation further enabled dual-color imaging of microglia and cortical vasculature over a large cortical area through the intact skull (Fig. 4i; Extended Data Fig. 3). These results collectively demonstrate that ClearGel provides sufficient clearing quality across the entire dorsal skull for cellular-resolution structural imaging.

Beyond whole-skull clearing, spatially selective multi-region clearing imposes additional demands on the clearing agent. With liquid-based skull optical clearing, freely diffusing reagents are typically confined only by a fixed chamber or contiguous liquid reservoir, rendering spatial control difficult to achieve without physical barriers. By contrast, in ClearGel, the extent and location of skull transparency can be precisely defined by the gel-covered area itself. We next tested whether optical access could be spatially specified simply by the application pattern of the gel. When ClearGel was applied only to discrete cortical targets, enhanced vascular visualization was strictly confined to those selected regions, whereas untreated areas remained at near-baseline levels (Fig. 4f). Likewise, hemispheric application selectively improved vascular contrast on one side of the skull without affecting the contralateral hemisphere (Fig. 4g). Thus, the transparent region was determined by where the gel was placed, rather than by a fixed chamber geometry or continuous liquid interface—a feature fundamentally distinct from existing liquid-based methods.

Together, these results establish spatial controllability as a defining advantage of ClearGel and demonstrate that the same gel-confined interface can provide high-quality transcranial optical access across multiple spatial scales, from focal cortical sites to hemispheric and whole-skull coverage, offering flexible experimental designs previously unattainable with conventional clearing approaches.

### ClearGel enables superior repeatable imaging over 14 days

Chronic transcranial imaging requires not only high-quality optical access at a single time point, but also the ability to reliably recover that access over repeated sessions. Previous studies have suggested that UV-cured window configurations are beneficial for long-term repeatable imaging^11,14^. To systematically evaluate ClearGel in this context, we compared two longitudinal imaging formats. In the reversible clearing mode (ClearGel-RC), the optical interface was re-established before each imaging session without permanent skull sealing. In the UV-cured window mode (ClearGel-UCW), a coverslip was permanently bonded to the skull after the initial clearing procedure, analogous to previously reported UV-based chronic window strategies^11,14^ (Fig. 5a, b).

**Fig 5.**
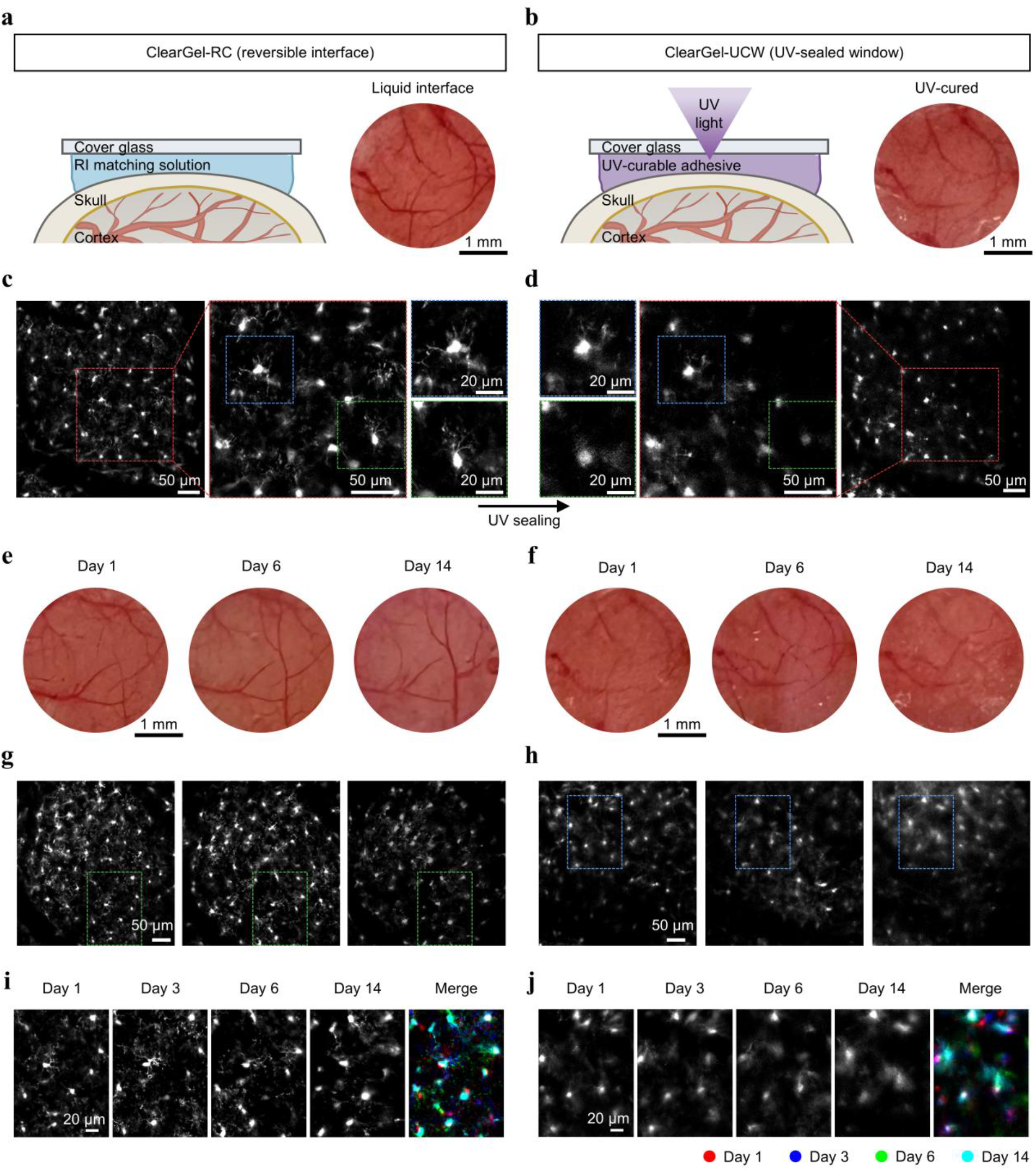
A reversible ClearGel interface supports repeated transcranial imaging over 14 days. **a, b**, Schematic of the reversible interface configuration (ClearGel-RC; **a**) and the UV-cured window configuration (ClearGel-UCW; **b**). In ClearGel-RC, refractive-index matching solution was placed beneath a removable coverslip to form a reversible liquid interface. In ClearGel-UCW, the coverslip was permanently bonded to the skull with a UV-curable adhesive after ClearGel treatment. The bright-field images in **a** and **b** show the same skull region before and after UV sealing, respectively. **c, d**, Paired two-photon 30 µm maximum-intensity projections of EGFP-expressing cortical microglia acquired from the same cortical field in the same 6-week-old *Cx*_*3*_*cr1*^*GFP/+*^ mouse before (**c**) and immediately after (**d**) UV sealing. Overview images, enlarged views of the boxed regions and higher-magnification views of selected microglia are shown. Color-matched boxes indicate the corresponding enlarged regions. **e, f**, Longitudinal bright-field images of the skull surface acquired on days 1, 6, and 14 using ClearGel-RC in a separate mouse (**e**) or ClearGel-UCW in the mouse shown in **a**-**d** (**f**). For ClearGel-RC, ClearGel treatment was repeated and the reversible liquid interface was re-established before each imaging session. **g, h**, Longitudinal two-photon 30 µm MIPs of the same cortical fields shown in **e** and **f**, respectively, acquired on days 1, 6 and 14. **i, j**, Higher-magnification 30 µm MIPs of the boxed regions in **g** and **h**, respectively, acquired on days 1, 3, 6, and 14 using ClearGel-RC (**i**) or ClearGel-UCW (**j**). Right, registered overlays showing the spatial correspondence of microglial somata across imaging sessions. Day 1, red; day 3, blue; day 6, green; day 14, cyan. **c, d, g**-**j**, 2 µm z-step, 0.25 µm per pixel. Experiments were independently replicated in n = 3 mice per configuration.

Two-photon imaging of cortical microglia in *Cx*_*3*_*cr1*^*GFP/+*^ mice revealed that ClearGel-RC consistently provided superior image quality compared with ClearGel-UCW. In the reversible format, fine microglial processes and cellular morphology were clearly resolved throughout the observation period. In contrast, imaging through the UV-cured window showed progressive degradation, with reduced contrast and loss of fine structural detail—a difference that was already evident immediately after UV curing and became more pronounced over repeated sessions from day 1 to day 14 (Fig. 5c–h). Quantitative analysis of the representative longitudinal images shown in Fig. 5g and h confirmed that ClearGel maintained higher signal-to-background ratio (SBR) and contrast-to-noise ratio (CNR) than ClearGel-UCW during repeated imaging (Extended Data Fig. 4).

We next examined whether ClearGel-RC could support repeated high-resolution access to the same cortical field across days. Using the reversible format, the same microglial population could be reliably revisited on days 1, 3, 6, and 14, with stable cellular positions and consistently resolved ramified morphology (Fig. 5i). By contrast, repeated imaging through the UV-cured window progressively lost cellular detail, making reliable high-resolution follow-up increasingly difficult (Fig. 5j). Together, these results demonstrate that ClearGel-RC enables chronic imaging through a reversible optical interface and that repeated re-clearing is more effective than permanent skull sealing for maintaining high-quality transcranial imaging over time.

To further validate the robustness of ClearGel-RC in pathological contexts, we adopted a laser-induced cortical injury model (Extended Data Fig. 5). Under this paradigm, ClearGel-RC enabled six consecutive days of repeated imaging without noticeable deterioration in image quality (Extended Data Fig. 5). After approximately 10 days, however, high-resolution imaging of the same cortical region could no longer be achieved. Importantly, this decline was attributable to chronic edema resulting from the laser injury^27-29^—a well-documented phenomenon that develops within days after cortical injury, persists for several weeks, and substantially alters local optical properties—rather than to any limitation of the clearing method itself. Thus, while ClearGel itself imposes no inherent constraint on longitudinal imaging, the natural progression of cortical pathology, such as post-injury edema, may ultimately limit optical access through any transcranial preparation.

### ClearGel minimally perturbs cortical glial homeostasis and supports deep vascular imaging in severely immunodeficient mice

To evaluate the biosafety and biocompatibility of ClearGel, we conducted a series of *in vivo* and *ex vivo* experiments to assess whether the technique induces cortical injury or immune responses. We first examined whether repeated ClearGel treatment perturbed cortical glial homeostasis. *In vivo* two-photon imaging of cortical microglia in *Cx*_*3*_*cr1*^*GFP/+*^ mice showed that the same microglial field remained morphologically stable before clearing, immediately after the first clearing round, 1 h later, and after a second clearing round performed 2 days later, with preserved soma positions and ramified processes throughout (Fig. 6a). *Ex vivo* imaging further revealed that microglial density and morphology in the cleared cortical region were comparable to those in the contralateral untreated cortex (Fig. 6b). Consistent with these observations, GFAP immunostaining showed no significant increase in astrocyte density after either a single ClearGel treatment or five repeated treatments, across both superficial cortical layers and the full cortical thickness, relative to untreated controls (Fig. 6c, d). H&E staining further revealed no apparent histological differences between the ClearGel-treated cortex and the contralateral control side (Extended Data Fig. 6). Collectively, these results indicate that repeated ClearGel treatment causes minimal detectable perturbation of cortical glial homeostasis.

**Fig 6.**
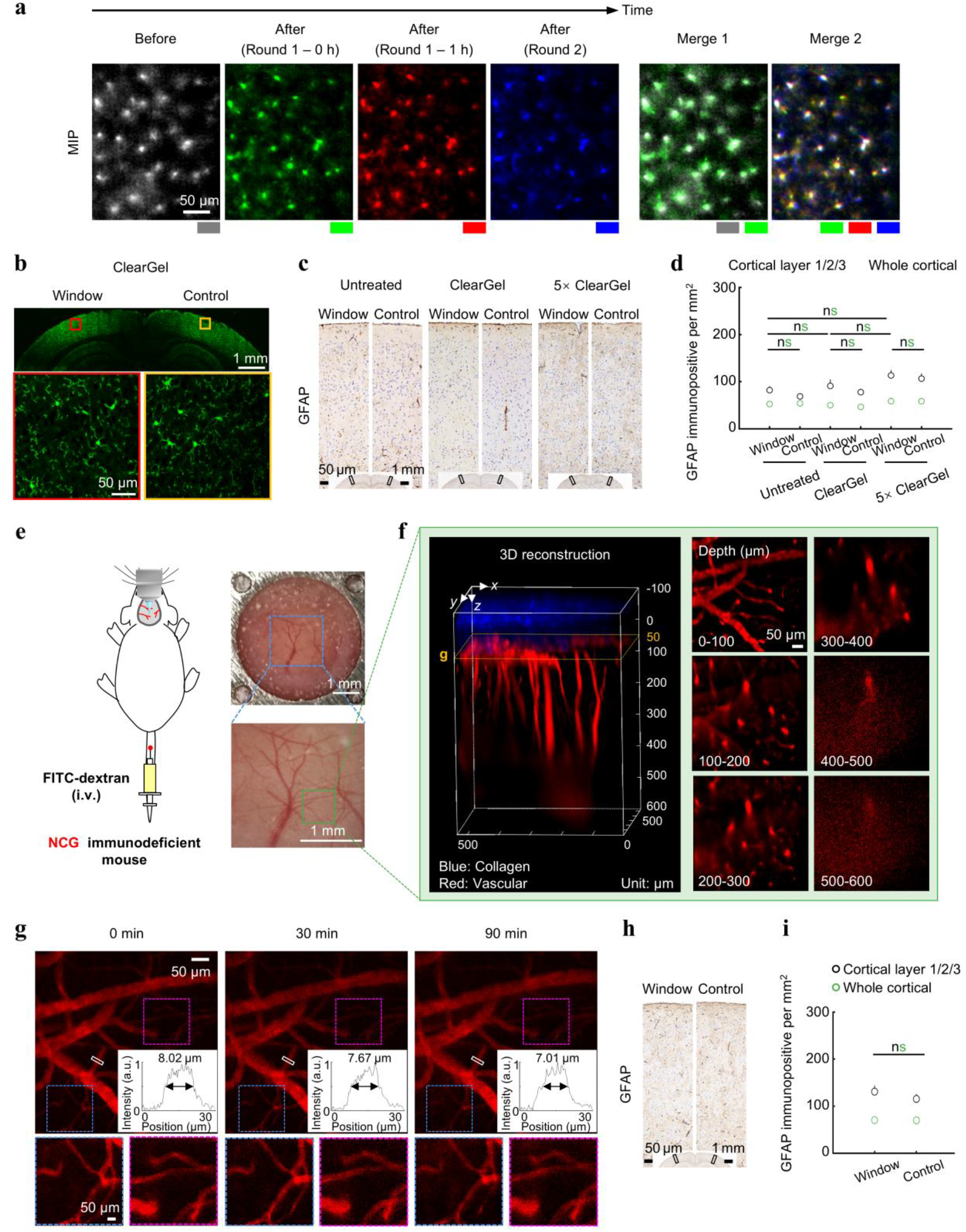
Repeated ClearGel treatment does not induce detectable cortical glial activation and enables deep transcranial vascular imaging in NCG mice. **a**, Longitudinal *in vivo* two-photon maximum-intensity projections (MIPs) of EGFP-expressing cortical microglia acquired from the same cortical field in a 6-week-old *Cx*_*3*_*cr1*^*GFP/+*^ mouse before ClearGel treatment, immediately after the first treatment (Round 1, 0 h), 1 h after the first treatment (Round 1, 1 h), and after a second treatment performed 2 d later (Round 2). Gray, before treatment; green, round 1 at 0 h; red, round 1 at 1 h; blue, round 2. Merge 1 overlays the before-treatment and round 1 - 0 h images; Merge 2 overlays the round 1 - 0 h, round 1 - 1 h and round 2 images. **b**, Representative *ex vivo* two-photon MIPs of cortical microglia in brain sections collected 2 d after ClearGel treatment. The cortex beneath the treated region (Window) and the matched contralateral cortex (Control) from the same section are shown. Bottom, enlarged views of the boxed regions. **c**, Representative GFAP immunostaining from untreated mice and from mice analysed 7 d after one ClearGel treatment or 14 d after five repeated treatments. The window-side and the matched contralateral control region are shown for each condition. **d**, Quantification of GFAP immunopositive cells per mm^2^ in cortical layers 1/2/3 and across the full cortical thickness for the conditions shown in **c**. Data are mean ± s.e.m.; n ≥ 3 mice. Data from cortical layers I–III and full cortical thickness were analyzed separately using one-way ANOVA followed by Tukey’s multiple-comparisons test. ns, not significant. **e**, Schematic of transcranial vascular imaging in 6-week-old female NCG mice after intravenous injection of 150 kDa FITC-dextran (10 mg ml^-1^). Right, representative bright-field image of the ClearGel-treated skull. The blue box in the upper image indicates the region enlarged below, and the green box indicates the two-photon imaging field shown in **f. f**, Three-dimensional two-photon reconstruction of cortical vasculature and skull collagen through the ClearGel-treated intact skull. Red, FITC–dextran-labeled vasculature; blue, collagen second-harmonic generation (SHG) signal. Right, depth-resolved MIPs acquired at the indicated depths to 600 µm below the pia. **g**, Representative two-photon images of the same cortical vascular field acquired at 0, 30, and 90 min. Boxed regions are enlarged below. Insets show fluorescence-intensity profiles across the indicated vessels; values denote the corresponding full width at half maximum (FWHMs). **h**, Representative GFAP immunostaining of NCG mouse cortex collected 7 d after vascular imaging. The cortex beneath the treated region (Window) and the matched contralateral cortex (Control) are shown. **i**, Quantification of GFAP immunopositive cells per mm^2^ in cortical layers 1/2/3 and across the full cortical thickness for the regions shown in **h**. Data are mean ± s.e.m.; n = 3-4 measurement sites from one representative mouse. Matched window and contralateral regions from the same mouse were compared using a two-tailed paired t-test. ns, not significant.

To further validate the biosafety of ClearGel in a more stringent context, we turned to a severely immunodeficient mouse model. NCG mice^30^ (GemPharmatech, t001475) lack functional T, B, and NK cells and are widely used in tumor and immune-related studies^31-34^, in which invasive cranial procedures are particularly undesirable^6^. We therefore tested whether ClearGel could provide minimally invasive optical access for cortical vascular imaging in this sensitive model. After intravenous FITC-dextran injection, ClearGel enabled three-dimensional transcranial vascular imaging to depths approaching 600 μm, resolving the cortical vascular hierarchy from large pial vessels to deeper penetrating vessels and capillaries (Fig. 6e, f). Image quality remained stable during 90 min of continuous imaging, as reflected by preserved capillary contrast and vessel-edge definition over time (Fig. 6g). To assess delayed astrocytic responses, NCG mice were examined 7 days after imaging; GFAP staining showed no significant difference in astrocyte density between the ClearGel-treated cortex and the contralateral control side (Fig. 6h, i). These findings extend the applicability of ClearGel to severely immunodeficient models, in which minimally invasive optical access is particularly advantageous for vascular, tumor, and neuroimmune studies.

## Discussion

In this study, we established ClearGel, a thermoresponsive hydrogel-based skull optical clearing strategy that enables simple, region-selective, and minimally invasive longitudinal transcranial brain imaging. We demonstrate that ClearGel supports spatially and temporally scalable imaging applications, including near-millimeter structural two-photon imaging, awake functional imaging of neuronal Ca^2+^ activity and cAMP dynamics, flexible optical access from focal sites to whole-skull coverage, repeated imaging over at least 14 days, and deep vascular imaging in severely immunodeficient mice.

A key innovation of ClearGel lies in the introduction of an OCA-loaded thermoresponsive hydrogel that transforms skull optical clearing from a freely diffusing liquid treatment into a gel-confined interfacial process. In the sol state, ClearGel can be applied to irregular skull surfaces and user-defined regions; upon contact with the warm skull, rapid in situ gelation creates a localized transparent layer that conforms to the skull surface while restricting lateral reagent diffusion. This confinement is particularly important near cranial sutures, where liquid reagents are prone to capillary spreading and local leakage^6,10,12,13^, potentially increasing the risk of vascular damage associated with locally high reagent concentrations. As a result, ClearGel enables region-selective clearing in focal, multi-site, hemispheric, and whole-skull formats without requiring a chamber or cap, while reducing uncontrolled reagent accumulation at vulnerable anatomical boundaries.

Compared with open-skull^5,9^ and thinned-skull^7,8^ preparations, ClearGel preserves skull integrity and avoids technically demanding procedures such as skull thinning, polishing, or craniotomy, thereby simplifying preparation and reducing the risk of mechanical or thermal damage to the meninges or cortex. Compared with conventional liquid-based skull clearing methods^6,10,12,13^, ClearGel defines the cleared region by the hydrogel-covered area, eliminating the need for a chamber, cap, or other liquid-retaining device. Importantly, this operational simplicity does not compromise optical performance: ClearGel significantly improved skull transparency and provided deeper transcranial TPM imaging than representative existing skull clearing methods. Unlike permanently sealed cleared-skull windows—in which a cured adhesive or polymer layer is applied after liquid skull clearing to maintain chronic transparency—ClearGel remains reversible and can be re-established before each imaging session, allowing re-clearing during longitudinal experiments. Notably, unlike UV-curable window strategies such as TIS^14^ and LCCW^11^, which rely on permanent resin sealing after an initial liquid clearing step and therefore cannot replenish the clearing agent or correct for gradual skull surface changes, ClearGel’s reversible format allows re-establishment of the optical interface before each imaging session. This distinction likely underlies the superior long-term image quality we observed with ClearGel-RC over UV-cured sealing. Thus, ClearGel is particularly suitable for studies requiring simple operation, flexible selection of cortical regions, deep transcranial imaging, and repeated optical access across days.

We also evaluated two possible strategies for longitudinal ClearGel-based imaging: reversible clearing (ClearGel-RC) and UV-cured sealing (ClearGel-UCW). Although both strategies maintained optical access, our results indicate that the reversible mode is preferable for preserving high-quality transcranial imaging over time. In the reversible mode, the effective RI of the cleared skulls (∼1.45) *in vivo* closely matched that of the ClearGel-S3 imaging solution (∼1.47), reducing RI discontinuities at the skull–reagent interface and helping to preserve image quality over time^6^. By contrast, the UV-curable resin exhibited a higher RI (∼1.51), which likely introduced a larger RI mismatch and contributed to the observed degradation in image performance^6^. Moreover, once the resin is cured, fresh clearing reagent can no longer be exchanged, and skull transparency may gradually decline as bone remodeling and surface changes proceed. In ClearGel’s reversible mode, optimal skull transparency can be re-established before each imaging session, offering both optical and practical advantages over the solidified window for longitudinal applications.

ClearGel also exhibited an excellent biosafety profile. Repeated ClearGel treatment did not induce obvious microglial activation, astrocytic response, or histological abnormalities, supporting the minimal invasiveness of the technique. This compatibility is likely aided by the gel-confined delivery mode: after *in situ* gelation, ClearGel remains localized on the skull surface, restricts lateral reagent spreading, and may reduce uncontrolled accumulation of clearing agents near cranial sutures or other vulnerable anatomical boundaries. Together with the diffusion barrier provided by the dura, this localized delivery likely limits perturbation of the underlying brain tissue^6^. Importantly, ClearGel also enabled deep transcranial vascular imaging in severely immunodeficient NCG mice, with stable image quality during continuous imaging and no detectable delayed astrocytic activation. These findings suggest that ClearGel can provide minimally invasive optical access even in immune-sensitive animals, rendering it potentially useful for longitudinal imaging in neuroimmune, tumor, disease, and humanized or immunodeficient mouse models.

Several aspects of ClearGel warrant further optimization. First, the current protocol requires an extended ClearGel-S1 incubation to establish a homogeneous transparent skull window, which may not be compatible with certain experimental designs requiring rapid optical access. Ultrasound-assisted transport or other permeability-enhancing strategies may accelerate reagent delivery while preserving gel confinement. Second, although ClearGel enables deep two-photon imaging, spatial resolution and contrast in deep cortical layers remain affected by residual scattering from the cleared skull and brain tissue, RI discontinuities among the objective immersion medium, S3 solution, cleared skull, and underlying brain tissue, and aberrations arising from the natural curvature and uneven surface of the skull. These effects may become more pronounced in older animals with thicker or more mineralized skulls. In 10-week-old mice, the maximum imaging depth decreased to approximately 650 μm (Extended Data Fig. 7). Further optimization of RI matching, together with adaptive optics, wavefront correction, or three-photon microscopy, may improve deep-layer imaging quality and extend the accessible imaging depth^35-37^. With these refinements, ClearGel may find broad applications in longitudinal studies of neural activity, neurovascular coupling, neuroimmune interactions, and disease progression, particularly in experimental settings where preservation of the intact skull environment is essential^38,39^.

## Acknowledgements

We thank Prof. Yang Zhan (Brain Cognition and Brain Disease Institute, Shenzhen Institutes of Advanced Technology, Chinese Academy of Sciences, Shenzhen, China) and Prof. Bo Peng (Institute for Translational Brain Research, Fudan University) for providing the Thy1-GFP-M and *Cx*_*3*_*cr1*^*GFP/+*^ mice, respectively. We also thank Prof. Yulong Li and Yu Zheng (School of Life Sciences, Peking University) for providing AAV9-hSyn-cAMPinG1 viruses.

## Author contributions

L. L. and W. Z. conceived and designed the study. L. L. developed and optimized ClearGel, performed the optical imaging experiments, analyzed the data and wrote the initial manuscript. W. Z. revised the manuscript. W. Z. and H. L. supervised the project. L. L., Y. G. and J. Y. performed wide-field two-photon microscopy imaging. Y. H. and N. X. provided code support. S. Y., L. W., and J. C. provided the cAMP virus and performed viral injections. T. W., J. L., L. J. L., C. L., and X. D. provided resources and technical advice.

